# *In situ*, High-Resolution Quantification of CO_2_ Uptake Rates via Automated Off-Gas Analysis Illuminates Carbon Uptake Dynamics in Cyanobacterial Cultures

**DOI:** 10.1101/2024.06.02.597031

**Authors:** Christopher M. Jones, Sean Innes, Steven Holland, Tyson Burch, Sydney Parrish, David R. Nielsen

**Author notes:** Corresponding author. BDC C499C, Tempe AZ 85282, United States.

## Abstract

Quantification of CO_2_ fixation rates by cyanobacteria is vital to determining their potential as industrial strains in a circular bioeconomy. Currently, however, CO_2_ fixation rates are most often determined through indirect and/or low-resolution methods, resulting in an incomplete picture of both dynamic behaviors and total carbon fixing potential. To address this, we developed a novel, low-cost system for *in situ* off-gas analysis which supports the automated acquisition of high-resolution and CO_2_ uptake rates from cyanobacterial cultures. Carbon fixation data obtained via this system was independently verified by elemental analysis of cultivated biomass. Using *Synechococcus sp.* PCC 7002 and *Synechocystis sp.* PCC 6803, we demonstrate that phototrophic CO_2_ uptake rates accelerate linearly to a maximum before then decaying monotonically to cessation by stationary phase. Furthermore, consistent with the expected stoichiometry, we found that strong correlations exist between both the rates and total levels of cell growth and carbon fixation. This system simultaneously provides a high-resolution growth curve, accurate carbon fixation rates, as well as the total amount of carbon fixed in a cyanobacterial batch culture, thus illuminating the parameters with which cyanobacterial researchers can exploit use to realize their full potential in industrial applications.

## 1. Introduction

Cyanobacteria are phototrophic microorganisms that represent a promising biotechnology platform for meeting carbon-neutral chemical demand. Most notably, cyanobacteria: i) derive their carbon and energy from CO_2_ and light, respectively; ii) do not require arable land nor fresh water for growth; iii) are easy to manipulate genetically; and iv) display higher aerial productivity rates than terrestrial plants (Dismukes, Carrieri, Bennette, Ananyev, & Posewitz, 2008; Krishnan et al., 2021; Melis, 2009). Key to benchmarking the performance of any bioproduction strain or system, however, is the ability to accurately quantify rates of substrate utilization with high resolution. Although a solved problem for most heterotrophic bioprocesses, straightforward and scalable methods do not yet exist for determining instantaneous rates of CO_2_ uptake in cyanobacterial cultures. Instead, the predominant focus remains on total levels of carbon uptake, as most commonly determined via correlation with cell growth using a biomass composition that is either assumed (e.g., cell dry weight is ∼50% carbon) or measured (via elemental analysis) (Aghaalipour, Güllü, & Akbulut, 2020; Moraes, da Rosa, Santos, & Costa, 2020; Oliver & Atsumi, 2015); a laborious and time-consuming approach that still only produces low-resolution datasets. Alternatively, instantaneous rates of CO_2_ uptake by phototrophic cultures have been estimated by measuring rates of O_2_ evolution (Benschop, Badger, & Dean Price, 2003; Price, Woodger, Badger, Howitt, & Tucker, 2004); also an indirect method whose results can be obscured by the occurrence of O_2_ consumption pathways. Direct measurements of instantaneous CO_2_ uptake rates can be obtained via H^14^CO ^-^ uptake experiments, though these must be performed *ex situ* and at the small scale (Schwarz, Friedberg, & Kaplan, 1988; Xu et al., 2008). Membrane inlet mass spectrometry (MIMS) has been used for the direct, *in situ* measurement of CO_2_ uptake rates by detecting changes in dissolved CO_2_ levels (Douchi et al., 2019; Leggat, Badger, & Yellowlees, 1999; van Hunnik, Amoroso, & Sultemeyer, 2002). Both direct methods, however, require access to more specialized analytical instrumentation.

As a more amenable approach to directly measuring rates of CO_2_ uptake, gas chromatography (Xiong et al., 2015) or an infrared gas analyzer (Doucha, Straka, & Lívanský, 2005; Oakley C.A., 2012) have instead been used to measure inlet and outlet gaseous CO_2_ concentrations (*C0_2(in)_*and *C0_2(out)_*, respectively), which together can be used to perform an overall ‘CO_2_ balance’ on the culture (Havlik, Lindner, Scheper, & Reardon, 2013). In this case, for a continuously bubbled batch culture, the volumetric CO_2_ uptake rate (*U*) is determined as the difference between the mass rate of gaseous CO_2_ entering vs. leaving the culture (Ohnishi et al., 2010), as reflected in Equation 1.

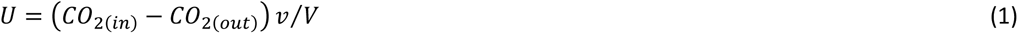

Where *v* is the volumetric flow rate of the gas stream (assumed to be equal for the inlet and outlet in a system with low backpressure) and *V* is the liquid volume of the culture. Note that, in a well buffered system maintained at constant pH, accumulated levels of dissolved inorganic carbon (C_i_) species remain constant and can be omitted from Equation 1. However, despite the straightforward implementation of this approach, current applications suffer from a need for *ex situ* analysis (Spalding & Ogren, 1985), low bandwidth, or are cost-prohibitive, especially if scaled in support of multiple cultures (Shabestary et al., 2021).

To address the limitations associated with the above methods, here we report on the development, validation, and use of an automated off-gas sampling system that supports the direct, *in situ* measurement of CO_2_ uptake rates with high-resolution from multiple cyanobacteria cultures in parallel. A central component of this low-cost and easy to assemble system is a custom designed, Arduino controlled mini-solenoid array that automatically directs inlet and outlet gas flows to a single infrared gas analyzer and in-line micro flow meter, thereby allowing instantaneous CO_2_ uptake rates to be measured across multiple cultures in a semi-continuous, high-resolution manner. Following its validation, this novel system was then used to illuminate, for the first time, the dynamic CO_2_ uptake profiles of two different cyanobacterial strains (*Synechococcus* sp. PCC 7002 and *Synechocystis* sp. PCC 6803) across a range of different light and CO_2_ regimes. This system represents a versatile tool for understanding the dynamic CO_2_ uptake rate behaviors of diverse cyanobacteria and algae, thereby facilitating metabolic engineering efforts for carbon-neutral chemical production.

## 2. Materials and Methods

### 2.1 Bacterial strains and growth conditions

*Synechococcus* sp. PCC 7002 was grown in MediaA+ medium. Precultures of PCC 7002 were grown at 10,000 ppm CO_2_ and 150 μE until reaching OD_730_ = 4-8. Cultures were then inoculated to an OD_730_ = 0.05 to start an experiment. *Synechocystis* sp. PCC 6803 was grown in BG-11 medium. Precultures of PCC 6803 were grown at 5,000 ppm CO_2_ and 50 μE until reaching OD_730_ = 5-7. Cultures were then inoculated to an OD_730_ = 0.2 to start an experiment. Antifoam 204 was included in all experiments at 0.002% (v/v). Following inoculation, all cultures were connected to the off-gas sampling system, the feed gas initiated at a constant flow rate between 40-80 mL/min, and the light intensity set to the desired level. Cell growth was measured as optical density at 730 nm (OD_730_), using a Biomate 3 spectrophotometer (Thermo), with dilutions performed as necessary to remain within the linear range (OD_730_ = 0.01-0.3).

### 2.2 Culture apparatus

Cultures were grown in 50 ml borosilicate glass tubes (24 x 150 mm) capped with 2-hole tapered rubber stoppers (size 4). An inlet tube for gas bubbling was constructed from a 1 mL glass pipette, with Teflon tape applied to ensure a tight seal between the tube and rubber stopper. The gas outlet port was created by inserting a polypropylene reducing barbed tubing adapter into the stopper then connecting a short segment of silicon tubing (1/4” I.D.) to the free end. An LED panel (110 x 22 cm) was constructed using a series of nine parallel LED strips (Samsung, CW5000K), with total light intensity being controlled via an independent power supply.

### 2.3 Design and operation of the off-gas sampling system

A circuit diagram of the off-gas sampling system is provided in **Supplemental Figure 1A**. Briefly, each individual solenoid circuit was constructed of an Arduino-activated Darlington transistor (Bridgold TIP120 TO-220 NPN Darlington Bipolar Power Transistor, 5A 60V HFE:1000, 3-Pin) where the output digital pin of the Arduino was connected to the base pin of the transistor via a 1K ohm resistor. The collector pin of the transistor was connected to the negative polarity of the solenoid (Aiyima 5V DC 2-Position 3-Way Electric Solenoid Valve) while the emitter pin of the transistor was connected to the ground of the Arduino. A 5V potential was supplied by the Arduino to the positive polarity of the solenoid via a diode snubber (Chanzon 1N4007 Rectifier Diode 1A 1000V DO-41). A total of 10 individual solenoids circuits were soldered in parallel to a circuit board, resulting in the complete autosampler system. Ten digital output pins on the Arduino were used to regulate the solenoids. An LED was included in each circuit to indicate which culture tube is being sampled at any given time. A foam board housing was constructed to contain the circuit board and Arduino microcontroller, with an operational window allowing for LED viewing and power supply connections. The inlet of each solenoid was connected to the off-gas from a corresponding culture tube and the outlet of each solenoid was connected to the collection manifold (gang valve) using PVC tubing (1/8” I.D. x ¼” O.D.). Off**-**gas flow rates were measured using a microflow sensor (Omron D6F-P0001A1), as controlled by a second, independent Arduino microcontroller. Calibration was performed using a mass flow controller to correlate flow rate with output voltage. The Arduino ‘serial monitor’ feature was used to record flow rates during all experiments. CO_2_ concentrations in both feed and off-gas samples were measured and recorded (at a frequency of one reading per min) using a Vaisala infrared CO_2_ gas meter with MI70 indicator and GMP70 probe. Custom scripts developed to control the autosampler and flow sensor are provided in the **Supplemental Text.** Detailed methods describing data collection, processing, and analysis are provided in the **Supplemental Text.**

### 2.4 Carbon analysis of biomass and supernatant samples

The final volume of triplicate cultures was determined and centrifuged at 3,800 x *g* for 20 min and the supernatants carefully removed. Cell pellets were frozen at -80°C then lyophilized for 24 h, after which dry cell weights were determined. Lyophilized biomass samples were analyzed with respect to their carbon content using a PE 2400 Series II CHN elemental analyzer (PerkinElmer). Total fixed carbon in each biomass sample was then determined by multiplying the measured percent carbon by the dry cell weight of the sample. Total dissolved carbon in supernatant samples was determined using a Shimadzu TOC-V analyzer and corrected by subtracting the total dissolved carbon measured in a MediaA+ blank.

## 3. Results and Discussion

### 3.1 Development and characterization of a CO_2_ off-gas sampling system for cyanobacterial cultures

To enable the semi-continuous measurement of volumetric CO_2_ uptake rates (*U*) across an array of cyanobacteria cultures, a custom-designed, automated off-gas sampling system was first constructed. The culture apparatus was comprised of a parallel array of 50 mL glass tubes (in this case, up to nine), each capped with a secure rubber stopper with inserted glass tubes for both gas sparging and venting. The vented off-gas from each tube was then directed towards a CO_2_ gas probe followed by a microflow sensor (**Figure 1**). Unique to this design, to allow for the sampling of multiple tubes while only requiring a single infrared gas analyzer (CO_2_ gas probe) and microflow sensor pair (thereby minimizing total costs), a parallel array of 2-position, 3-way mini-solenoids was assembled, with a dedicated solenoid connected to the off-gas stream of each tube. The solenoid array was controlled using an Arduino microcontroller and PC, with a custom code developed to automate systematic toggling between and activation of each solenoid (see **Supplemental Figure 1** for a solenoid circuit diagram and **Supplemental Text** for the Arduino code). In the absence of activation, the off-gas from each culture exits unimpeded out of the rear of the solenoid whereas, when activated, the solenoid instead diverts it through a collection manifold and ultimately to the infrared gas analyzer and microflow sensor pair.

**Figure 1.**
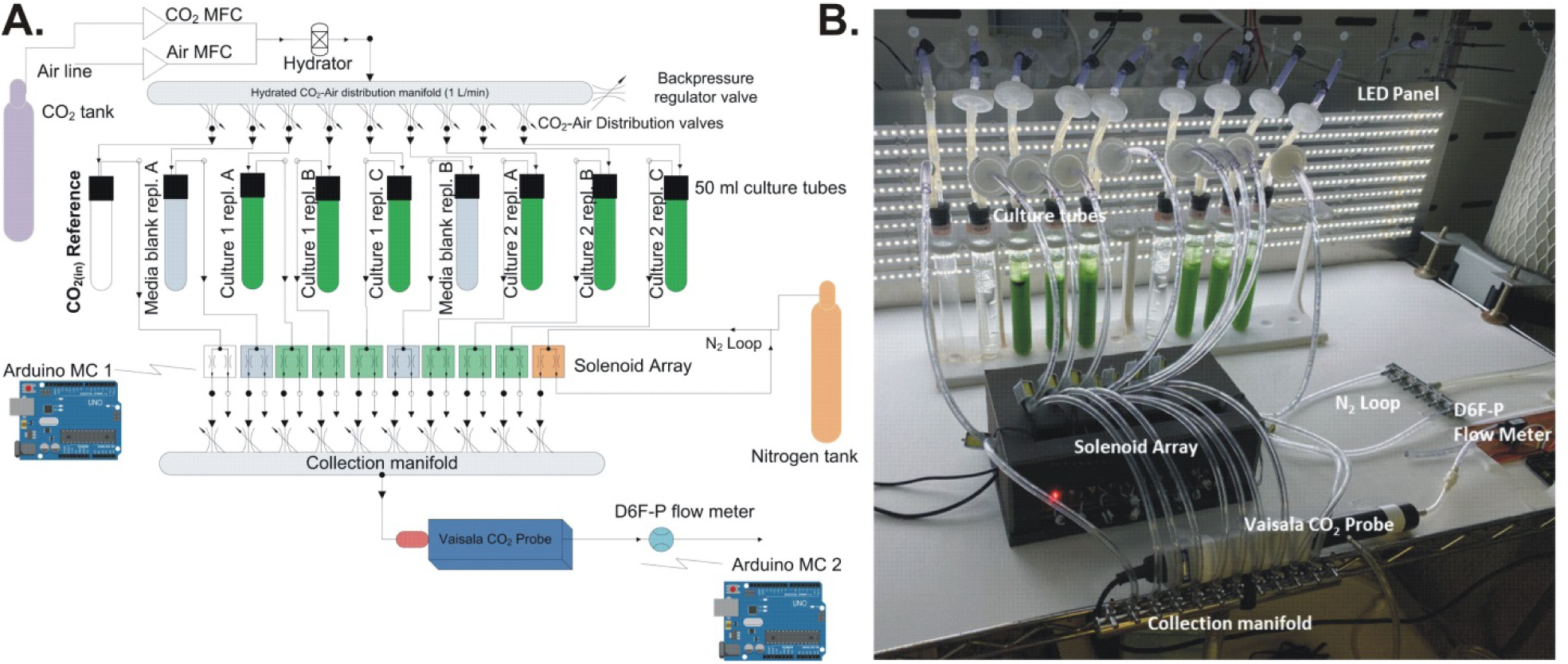
Overview of the automated CO_2_ off-gas sampling system. **A)** System schematic. A CO_2_-air feed gas mixture is established using mass flow controllers (MFCs) then hydrated and sent to a gang valve manifold to split the feed gas and control the flow rate to each individual culture tube. An empty tube serves as the *CO_2(in)_* reference while two media blanks are used to correct for background CO_2_ removal via absorption. The remaining tubes are used for two triplicate sets of cell culture samples. The off-gas leaving each tube enters a dedicated 3-way, two position mini-solenoid. In the absence of activation, the off-gas exits to the rear of the mini-solenoid. Upon activation, the off-gas is diverted towards a collection manifold connected to an infrared gas analyzer and microflow sensor. The mini-solenoid sampling program is controlled by an Arduino microcontroller (MC) according to a custom script. Each tube is sampled continuously for 15 minutes. Between tubes, N_2_ is used to purge the system for 4 minutes, which further aids in delineating between datasets. **B)** Photograph of the system in operation.

In a typical experiment, a prescribed CO_2_-air mixture (i.e., *C0_2(in)_* at flow rate *v*) is first established using a pair of mass flow controllers, then hydrated by bubbling through deionized water. The hydrated feed gas is then distributed among each of the cultures via a gang valve manifold, from which the flow rate to each tube is then controlled (here, between 40-80 mL/min). In addition to the set of six culture tubes (e.g., two sets of three replicates), also included is an empty tube (serving as a reference sample for *C0_2(in)_*) and two blank media tubes to correct for background CO_2_ removal via absorption alone. Following inoculation, each tube is then automatically sampled via the successive activation of its associated solenoid. Between tube-to-tube transitions, pure N_2_ gas (whose flow is controlled by another, independent solenoid) is used to purge the off-gas lines including the infrared gas analyzer and microflow sensor. To set an appropriate sampling time for each tube, the minimum equilibration time for the infrared gas analyzer was first empirically determined to be approximately 10 min at a flow rate of 40-80 mL/min (**Supplemental Figure 2**). Accordingly, each tube was continuously sampled for 15 min, with the signal averaged values obtained during the final 5 min serving as the final recorded values of CO_2_ concentration and flow rate. In particular, both CO_2_ concentration and flow rate measurements are subject to a high degree of variance; the likes of which is inversely proportional to the inlet conditions (**Supplemental Figure 2G,H**). For example, a coefficient of variation for the flow rate of approximately 10-20% was consistently observed (**Supplemental Figure 2H**). Nevertheless, by accounting for such factors and through repeated design-build-test cycles, we ultimately found that high-resolution, instantaneous measurements of CO_2_ uptake rates could be achieved for two independent cyanobacterial cultures (each in triplicate). Under such conditions, and based on the above equilibration constraints, CO_2_ and flow rate measurements were obtained for each tube at a frequency of once every 3 hours, or 8 samples/day.

Once fabricated, the off-gas sampling system was first applied to investigate the dynamic CO_2_ uptake rate behavior of wild-type PCC 7002 when grown under continuous illumination at a light intensity of 150 μE and inlet CO_2_ concentration set point of 5000 ppm (5048 ± 45 ppm over the course of the experiment). **Figure 2A** and **2B** respectively display an example of the raw CO_2_ and flow rate measurements obtained across all sample tubes during a total of eight consecutive loops (a 24 hour period) of the autosampler routine. From these raw data, volumetric rates of CO_2_ entering (**Figure 2C**) and exiting (**Figure 2D**) each culture tube can be instantaneously determined, their difference being the volumetric CO_2_ uptake rate (*U*; **Figure 2E**). Here, the volumetric CO_2_ uptake rate accelerates linearly until peaking at 1.70 ± 0.04 g CO_2_/L_culture_/day by 1.76 ± 0.13 days. By integrating the dynamic CO_2_ uptake rate data across the duration of the experiment, cumulative fixed carbon per volume of culture is then also revealed (**Figure 2F**). In this case, cumulative fixed carbon reached 1.93 ± 0.11 g carbon/L_culture_ after 9 days of continuous cultivation. Furthermore, the dynamics of cumulative fixed carbon correlate closely with those of cell growth (**Figure 2F**), as would be expected for phototrophic growth.

**Figure 2.**
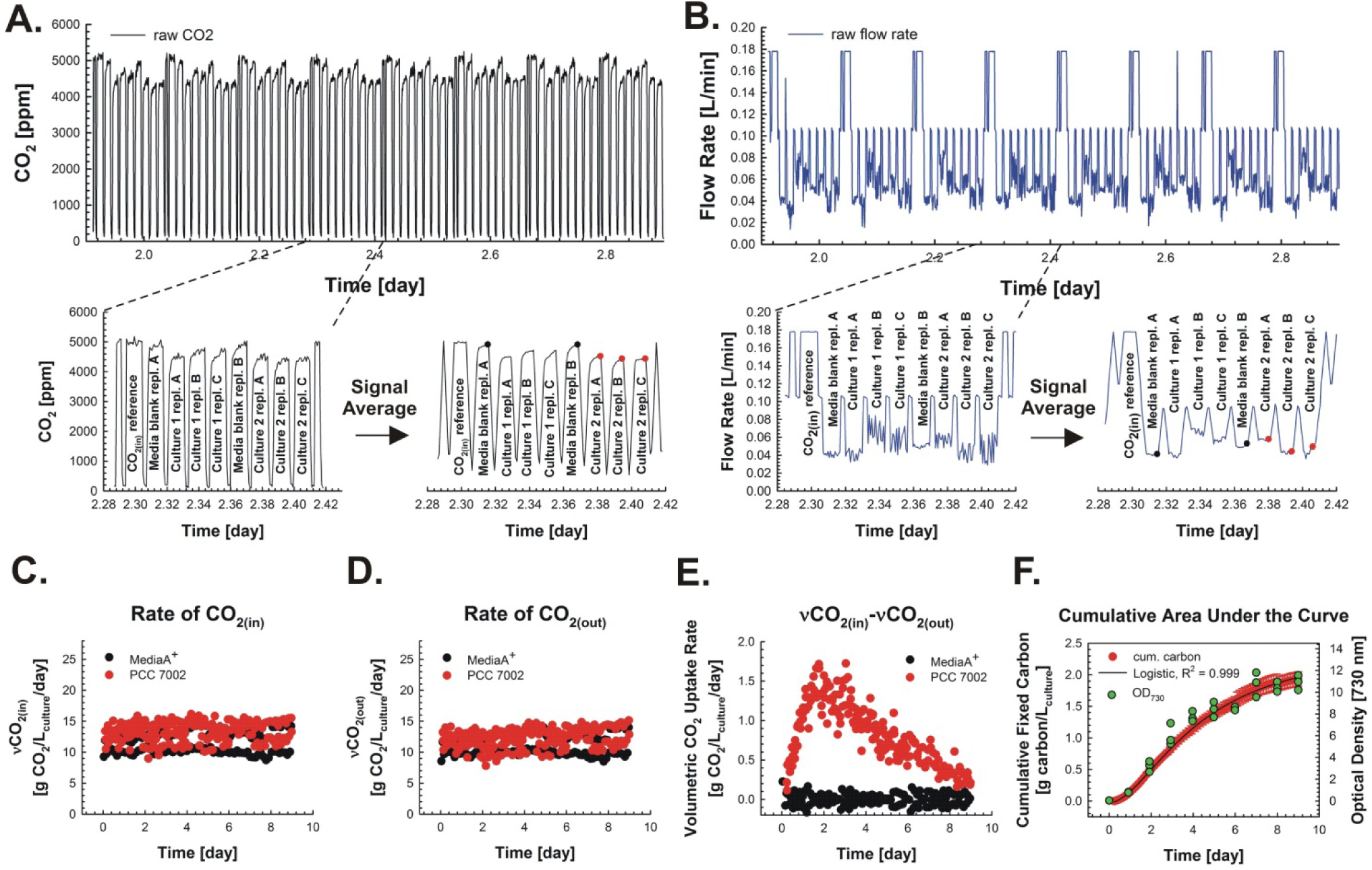
Data collection and processing workflow for the automated CO_2_ off-gas sampling system. **A)** An example of the raw data collected by the CO_2_ probe over a single (24 h) sampling period. The autosampler cycles through samples in a continuous loop as follows: *C0_2(in)_* reference, media blank-replicate 1, culture 1-replicate A, culture 1-replicate B, culture 1-replicate C, media blank-replicate 2, culture 2-replicate A, culture 2-replicate B, culture 2-replicate C. **B)** An example of the raw data collected by the inline flow sensor over a single (24 h) sampling period. Magnified regions in A) and B) show the corresponding raw data collected from each culture tube (individually labelled as indicated) during a single sampling loop (3 h) as well subsequent signal averaged data where colored dots correspond to those values obtained during the final 5 min of sampling which are then used in downstream data processing steps. **C)** Mass flow rate of CO_2_ entering the media blank (black) and culture tubes (red; one set of culture replicates shown) per volume of culture. **D)** Mass flow rate of CO_2_ exiting the media blank (black) and culture tubes (red; one set of culture replicates shown) per volume of culture. **E)** Volumetric CO_2_ uptake rate, determined as the difference between the mass flow rate of CO_2_ entering and exiting the media blank (black) and culture tubes (red; one set of culture replicates shown). To account for chemical absorption alone, averaged data from the media blanks at each time point is further subtracted from both data sets at each time point. **F)** Cumulative fixed carbon per volume of culture determined by integrating the data in E) using the point-slope method. Superimposed is cell growth (OD_730_). Data were generated from three independent cultures of wild-type *Synechococcus* sp. PCC 7002 grown under continuous illumination at 150 μE and feed gas concentration of 5,000 ppm (0.5%) CO_2_.

### 3.2 Quantifying and validating levels of cumulative fixed carbon

The ability of the off-gas sampling system to accurately quantify carbon fixation performance was next validated by comparing autosampler-determined levels of cumulative fixed carbon with direct measurements obtained via elemental analysis of produced biomass. Towards this end, PCC 7002 was grown under continuous illumination at 300 μE and inlet CO_2_ concentration set point of 10,000 ppm (1%; 10217 ± 126 ppm). As seen in **Figure 3**, the instantaneous CO_2_ uptake rate again accelerates linearly before peaking in this case at a maximum of 4.13 ± 0.14 g CO_2_/L_culture_/day within the first 24 hours then decelerating monotonically across the next 3 days. Indeed, the measured CO_2_ uptake rate became statistically insignificant relative to a blank MediaA+ sample by day 4 (**Figure 3A**), the point at which the culture also reached stationary phase (**Figure 3B**). Meanwhile, the dynamic behavior of the cumulative fixed carbon profile also closely reflects that of biomass growth; the former of which is well fit to a logistic function, with the rise of the curve representing the so-called ‘linear growth phase’ (Schuurmans, Matthijs, & Hellingwerf, 2017) (**Figure 3B**). Lastly, end-point samples of both the lyophilized cell pellet and clarified supernatant were analyzed with respect to their total carbon content and together used to provide a conventional determination of total carbon fixation by the culture. As seen in **Figure 3C**, this amount was statistically insignificant relative to the cumulative fixed carbon value obtained using our off-gas sampling system (**Figure 3C**). An analogous result was also obtained when the same experiment was repeated under lower light and CO_2_ conditions (150 μE and 0.25% CO_2_; **Supplemental Figure 3**), further validating the capabilities of the system. Thus, like elemental analysis of biomass, this novel method is similarly capable of accurately determining total carbon fixation, but does so in a way that also uniquely enables volumetric CO_2_ uptake rates to be instantaneously resolved in a dynamic and high-resolution manner.

**Figure 3.**
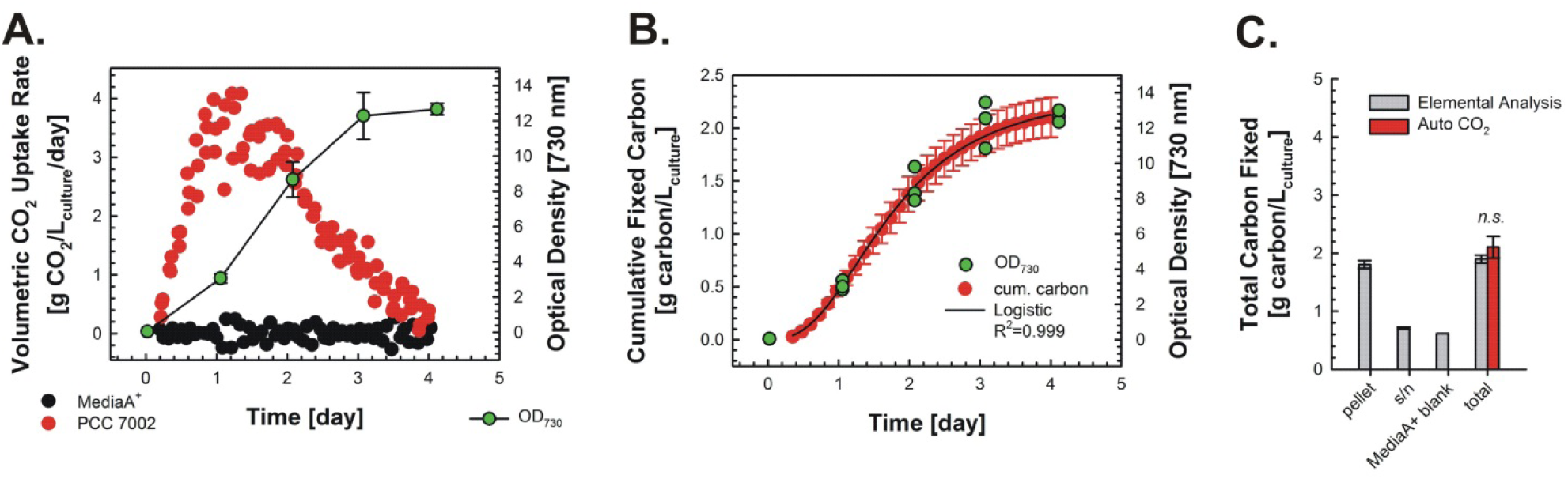
Comparing estimates of carbon fixation obtained using the CO_2_ off-gas sampling system with biomass elemental analysis. Wild-type *Synechococcus* sp. PCC 7002 was grown in MediaA+ under continuous illumination at 300 μE and a feed gas containing 10,000 ppm (1%) CO_2_. **A)** Volumetric CO_2_ uptake rate and cell growth (OD_730_) determined instantaneously during batch culture. **B)** Cumulative fixed carbon per volume of culture determined by integrating the data in A) using the point-slope method and further fitted via a logistic function (with *k* = 2.389 g carbon/L_culture_) using Sigma Plot v.11. Superimposed is cell growth (OD_730_). **C)** Comparing total levels of carbon fixation by the culture as predicted via the off-gas sampling system versus measured via biomass elemental analysis. Total carbon analysis was performed via elemental analysis of the lyophilized cell pellet harvested at the end of the culture (‘pellet’). Total carbon analysis was performed on the supernatant (‘s/n’) and a MediaA+ blank via TOC analysis. ’Total’ was determined by a summation of the carbon content of both lyophilized cell pellets and media samples as described in the methods. Error bars represent the standard deviation from biological triplicates. *n.s.* is not statistically significant (*p = 0.151*, Student’s t-test, normally distributed, equal variance).

### 3.3 Investigating the influence of CO_2_ delivery rate on PCC 7002 growth and CO_2_ fixation

With the system validated, we next sought to use it to explore the effects of CO_2_ delivery rate on growth and CO_2_ fixation by PCC 7002. Since the CO_2_ delivery rate can be influenced by the feed gas flow rate as well as its CO_2_ concentration, both were investigated, beginning with the former. PCC 7002 was grown under continuous light at 300 μE and inlet CO_2_ concentration set point of 10,000 ppm (1%) but two different flow rates: ‘low flow’, representing 1 volume gas per min per volume culture (1 vvm), and ‘high flow’, representing 2 vvm (corresponding to 45 ± 5 and 100 ± 4 mL/min, respectively, for a 50 mL culture volume); conditions supporting a statistically significant difference in the mean CO_2_ delivery rate over the course of the experiment (20.1 ± 1.3 vs. 44.3 ± 2.7 g CO2 /L_culture_/day; *p<0.001*, Mann-Whitney Rank Sum Test) (**Figure 4A**). The higher CO_2_ mass delivery rate expectedly resulted in a faster initial growth rate (**Figure 4B**) and, consistent with that, a higher maximum volumetric CO_2_ uptake rate (3.5 ± 0.4 vs. 2.7 ± 0.1 g CO_2_/L_culture_/day; **Figure 4C,D**).

**Figure 4.**
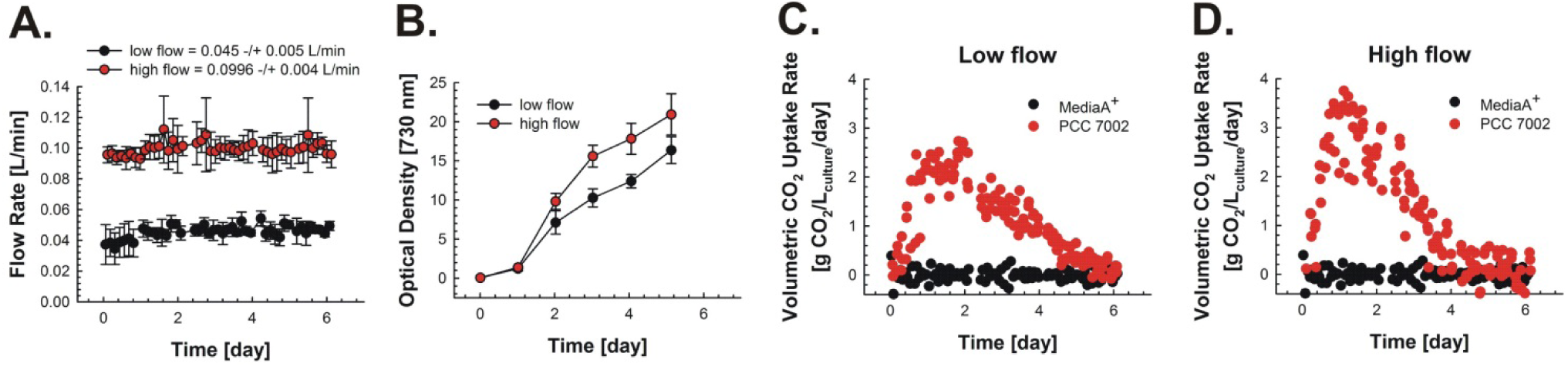
Investigating the effects of CO_2_ delivery rate on the rates and levels of CO_2_ fixation and growth by varying gas flow rate. Wild-type *Synechococcus* sp. PCC 7002 was grown in MediaA+ under continuous illumination at 300 μE and a feed gas containing 10,000 ppm (1%) CO_2_ at two different flow rates. **A)** Average flow rates of the two conditions, where ‘low flow’ and ‘high flow’ were approximately 1 and 2 vvm, respectively. **B)** Cell growth (OD_730_). **C)** Volumetric CO_2_ uptake rate determined instantaneously during batch culture under ‘low flow’ conditions. **D)** Volumetric CO_2_ uptake rate determined instantaneously during batch culture under ‘high flow’ conditions. Error bars represent the standard deviation from biological triplicates.

The effects of increasing the CO_2_ delivery rate by changing the feed gas concentration (*C0_2(in)_*) were likewise next examined. Here, PCC 7002 was grown under continuous light at 75 or 150 μE and inlet CO_2_ concentration set point between 1,000 to 7,500 ppm. As seen in **Figure 5A**, increasing *C0_2(in)_* also expectedly increased the maximum volumetric CO_2_ uptake rate at both light intensities. Moreover, the maximum volumetric CO_2_ uptake rate ultimately displayed light-dependent saturation kinetics with respect to *C0_2(in)_*, approaching approximately 2 g CO_2_/L_culture_/day at 150 μE and 1.6 g CO_2_/L_culture_/day at 75 μE (**Supplemental Figure 4**). Meanwhile, as *C0_2(in)_* was increased, the maximum volumetric CO_2_ uptake rate also occurred earlier in the culture (e.g., about day 9 at 1,000 ppm vs. about day 2 at 5,000 ppm) whereas the overall duration of active CO_2_ uptake became compressed, consistent with a faster growth rate. Indeed, levels of cumulative fixed carbon as well as rates and levels of cell growth were also higher as *C0_2(in)_* was increased (**Figure 5B**). Interestingly, acceleration of the volumetric CO_2_ uptake rate (i.e., initial slope of the volumetric CO_2_ uptake rate curve) also increased with increasing light intensity (**Figure 5A**), indicating that initial growth acceleration is more sensitive to light than carbon availability. Later in the growth phase, however, the opposite is true. Notably, the collective outcomes here are consistent with those of studies involving other phototrophs, where increasing *C0_2(in)_* has similarly resulted in increased CO_2_ uptake rates by *Synechocystis* sp. PCC 6803 (McGinn, Price, Maleszka, & Badger, 2003), *Synechococcus* sp. PCC 7942 (Omata et al., 1999), *Scenedesmus obliquus* (Azov, 1982), *Chlamydomonas reinhardtii* (Palmqvist, Sjoberg, & Samuelsson, 1988), as well as species of red algae (Johnston, Maberly, & Raven, 1992). Meanwhile, however, as also noted by others (Pruvost, 2022), despite supporting greater maximum volumetric CO_2_ uptake rates, increasing *C0_2(in)_* also unfortunately led to a decrease in CO_2_ utilization efficiency by the culture, which occurs as a result of mass transfer limitations and disparate rates of supply vs. fixation. Specifically, for example, when PCC 7002 was grown under continuous illumination at either 75 or 150 μE, the maximal CO_2_ utilization efficiency decreased from about 25% at a *C0_2(in)_* of 1,000 ppm to just about 8% at 7,500 ppm (**Supplemental Figure 5).**

**Figure 5.**
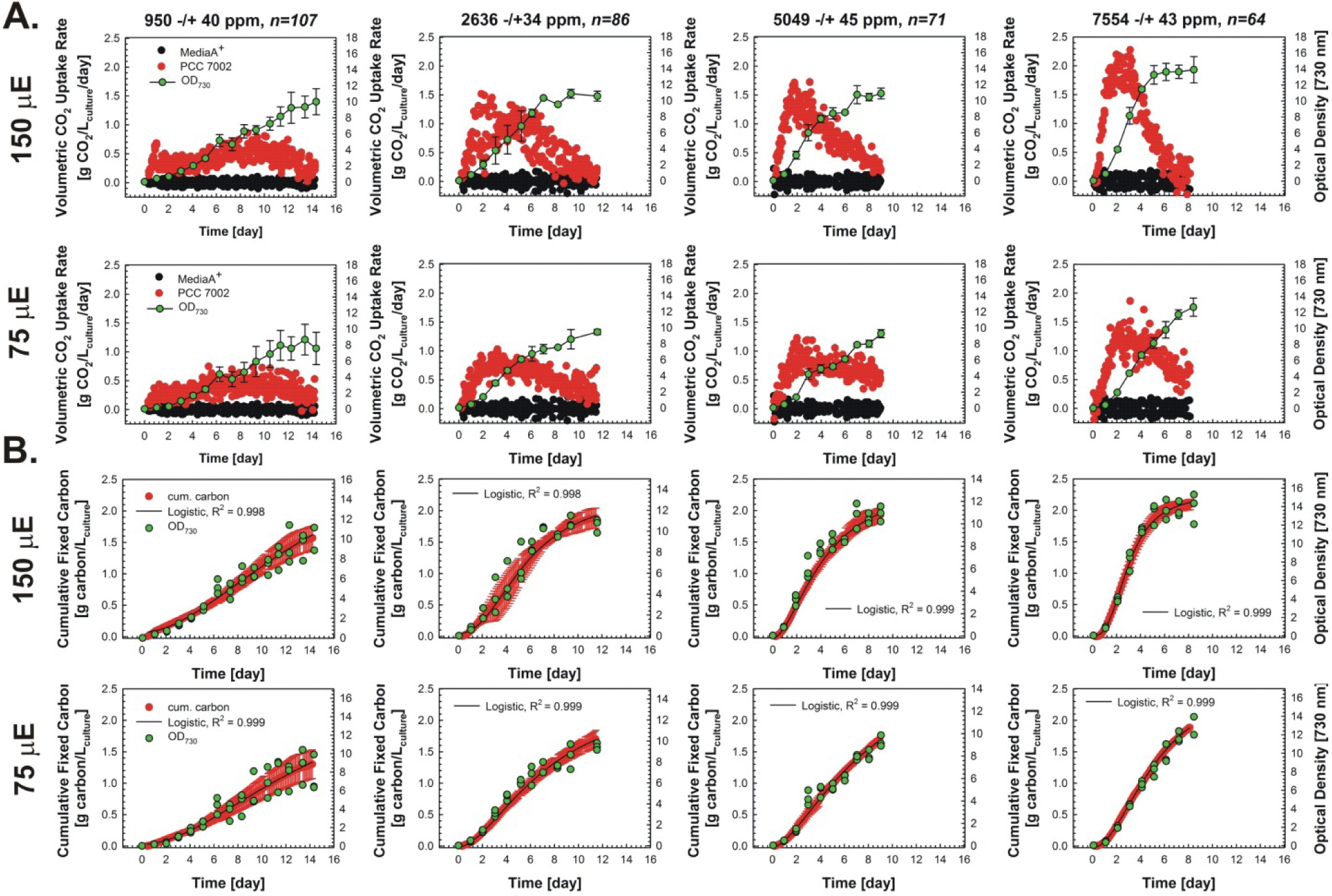
Investigating the effects of CO_2_ delivery rate on the rates and levels of CO_2_ fixation and growth by varying *CO_2(in)_* under different light regimes. Wild-type *Synechococcus* sp. PCC 7002 was grown in MediaA+ under continuous illumination at 75 or 150 μE and a feed gas containing 1,000, 2,500, 5,000, or 7,500 ppm CO_2_ (empirically determined mean values and standard deviations determined over the course of each experiment are presented in the respective figure headers) **A)** Volumetric CO_2_ uptake rate determined instantaneously during batch culture as a function of light intensity and *CO_2(in)_*. **B)** Cumulative fixed carbon per volume culture at 75 and 150 μE. Superimposed is cell growth (OD_730_). Error bars represent the standard deviation from biological triplicates.

### 3.4 Investigating the influence of light intensity on PCC 7002 growth and CO_2_ fixation

We next sought to examine the effects of light intensity on growth and CO_2_ fixation by PCC 7002. Here, the feed gas composition and delivery rate were held constant (7,500 ppm and 61 ± 4 mL/min) while increasing the intensity of continuous light; specifically, 75, 150, 300, and 600 μE. As with increasing *C0_2(in)_*, increasing light intensity expectedly resulted in greater maximum CO_2_ uptake rates while the overall duration of active CO_2_ uptake again became compressed (**Figure 6A**). The maximum CO_2_ uptake rate also displayed saturation kinetics with respect to light intensity, approaching a peak value of approximately 3 g CO_2_/L_culture_/day under the conditions examined (**Figure 6C**). The optical density-based growth curves and volumetric CO_2_ uptake rate profiles were fit to logistic and Weibull functions, respectively, and plotted on the same graph to more clearly demonstrate their direct relationship to light intensity, both of which increased with increasing light intensity (**Figure 6D,E**). Interestingly, while faster initial growth rates and greater cumulative fixed carbon were observed as light intensity was increased from 75 to 300 μE, by 600 μE, no further growth rate increase was observed and cumulative fixed carbon was significantly reduced relative to lower light intensities (**Figure 6B**). Since the total biomass yield at 600 μE was not significantly different than at 300 μE, this suggests that the specific carbon fixation rate (i.e., per cell) was reduced at the high light condition. This prospect was illuminated by normalizing the cumulative fixed carbon by the overall biomass yield (i.e., final OD_730_) which indeed revealed a decrease in total fixed carbon per biomass as light intensity was increased above 300 μE (**Figure 6F**); an outcome suggesting that carbon fixation was subject to photoinhibition under these conditions. Whole cell UV-Vis scans performed on day 2 of both the 300 and 600 μE cultures indicated a significant decrease in both chlorophyll and phycobilisome content at 600 μE compared to 300 μE (**Supplemental Figure 6**), further corroborating that reduced carbon fixation under these conditions was likely caused by high-light stress.

**Figure 6.**
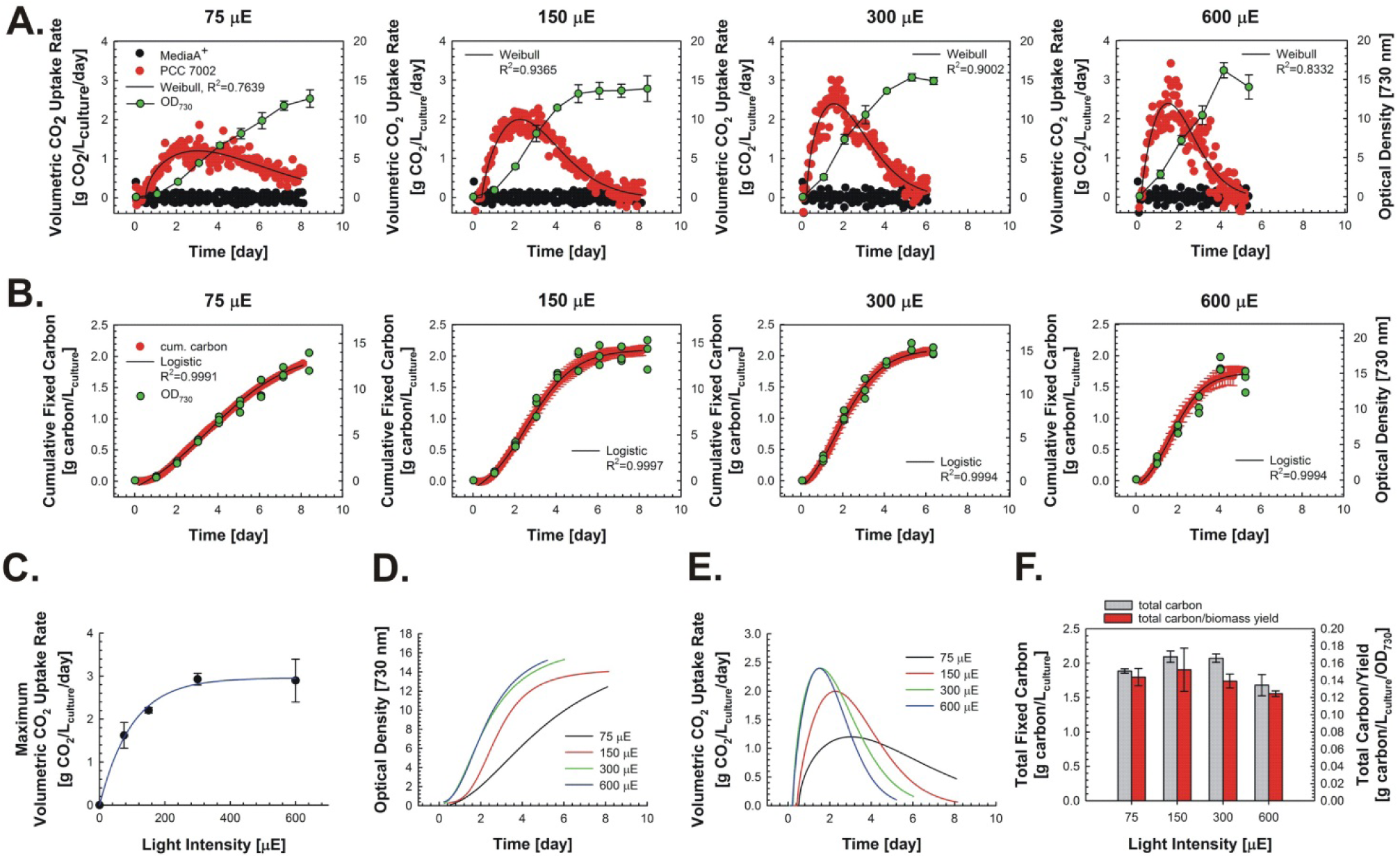
Investigating the effects of light intensity on the rates and levels of CO_2_ fixation and growth by PCC 7002. Wild-type *Synechococcus* sp. PCC 7002 was grown under continuous illumination at the indicated light intensities and 7,500 ppm (0.75%) CO_2_. **A)** Volumetric CO_2_ uptake rate determined instantaneously during batch culture as a function of light intensity. Solid line represents the best fit curve for a 5-parameter Weibull function. Superimposed is cell growth (OD_730_). **B)** Cumulative fixed carbon per volume of culture determined by integrating the data in A.) using the point-slope method and further fitted via a logistic function sing Sigma Plot v.11. Superimposed is cell growth (OD_730_). **C)** Maximum volumetric CO_2_ uptake rate at as a function of light intensity. **D)** The logistic fits of the growth rates as different light intensity. **E)** Weibull fits of the CO2 uptake rates at different light intensities. **F)** Total fixed carbon and total fixed carbon per biomass yield. Error bars represent the standard deviation from biological triplicates.

### 3.5 Investigating the influence of light intensity on growth and CO_2_ fixation by PCC 6803

To demonstrate the broader utility of the off-gas sampling system, we next applied it to elucidate the CO_2_ uptake rate profile of PCC 6803 and the effects of light intensity. PCC 6803 was cultured in BG-11 using a feed gas (64 ± 4 mL/min) with an inlet CO_2_ concentration set point of 5,000 ppm (0.5%) and four continuous light regimes: 50, 100, 200, and 400 μE. Like PCC 7002, we observed a proportional increase in volumetric CO_2_ uptake rate as light intensity was increased (**Figure 7A**). And, while initial cell growth rate increased between 50 and 100 μE, overall growth was reduced at and above 200 μE (**Figure 7A,D**). Different than PCC 7002, here these contrasting results suggest that the specific carbon fixation rate was instead elevated under high light conditions. Indeed, while the total amount of fixed carbon was not significantly different among groups (One-way ANOVA, n=3), total fixed carbon per biomass yield was markedly higher at 200 and 400 μE compared to lower light conditions (**Figure 7F**). This suggests that, while high light negatively impacts PCC 6803 growth, it has a positive effect on CO_2_ uptake. We suspect that additional carbon fixed at high light intensities may be being diverted to carbon storage compounds (e.g., polyhydroxybutyrate (PHB) and/or glycogen), as has been previously demonstrated in PCC 6803 (Monshupanee & Incharoensakdi, 2014).

**Figure 7.**
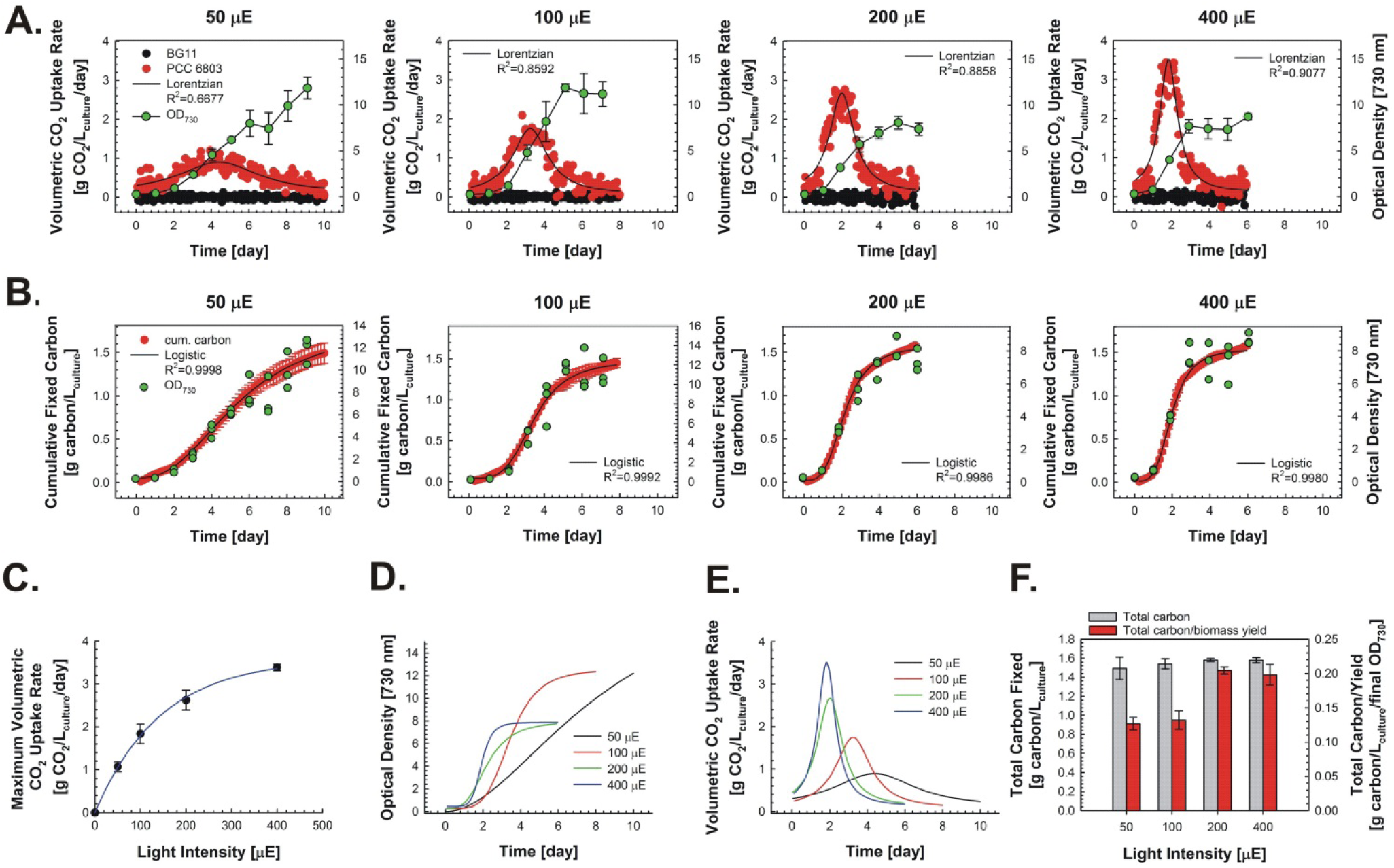
Investigating the effects of light intensity on the rates and levels of CO_2_ fixation and growth by PCC 6803. Wild-type *Synechocystis* sp. PCC 6803 was grown under continuous illumination at the indicated light intensities and 5,000 ppm (0.5%) CO_2_. **A)** Volumetric CO_2_ uptake rate determined instantaneously during batch culture as a function of light intensity. For 100, 200, and 400 μE, cultures began at 50 μE and were switched to the indicated light intensity on day 1. Solid line represents the best fit curve for a 4-parameter Lorentzian function. Superimposed is cell growth (OD_730_). **B)** Cumulative fixed carbon per volume of culture determined by integrating the data in A.) using the point-slope method and further fitted via a logistic function using Sigma Plot v.11. Superimposed is cell growth (OD_730_). **C)** Maximum volumetric CO_2_ uptake rate at as a function of light intensity. **D)** The logistic fits of the growth rates as different light intensity. **E)** Lorentzian fits of the CO2 uptake rates at different light intensities. **F)** Total fixed carbon and total fixed carbon per biomass yield. Error bars represent the standard deviation from biological triplicates.

### 3.6 Using high-resolution carbon fixation data to illuminate cyanobacterial growth kinetics

Throughout this study, the strong correlations observed between the profiles of cell growth and cumulative fixed carbon suggest that a direct proportionality should also exist between instantaneous measurements of growth and volumetric CO_2_ uptake rates. To illuminate this concept, we analyzed three independent data sets obtained using the off-gas sampling system. The first consisted of PCC 6803 grown at 5,000 ppm (0.5%) CO_2_ and continuous illumination at 150 μE. As seen in **Figure 8A**, following a protracted lag phase, cell growth began, leading to an increasing volumetric CO_2_ uptake rate which ultimately approached a maximum of 2.8 ± 0.3 g CO_2_/L_culture_/day by about day 5. Again, the profiles of cell growth and cumulative fixed carbon were strongly correlated (**Figure 8B**). And, as seen in **Figure 8C**, instantaneous estimates of volumetric growth rate (i.e., derivative of OD_730_ vs. time) were indeed strongly correlated with volumetric CO_2_ uptake rate. The same analysis was next performed for PCC 7002 grown under air and continuous illumination at 250 μE. Under these conditions, strict linear growth of PCC 7002 was observed along with a near constant, albeit low, volumetric CO_2_ uptake rate (**Figure 8D**). Strong correlations were again observed between both cell growth and cumulative fixed carbon (**Figure 8E**) as well as volumetric cell growth and CO_2_ uptake rates (**Figure 8F**). Lastly, the same analysis was also performed for PCC 7002 grown under 7,500 ppm CO_2_ and continuous illumination at 150 μE (data from **Figure 5**). Under these conditions, the volumetric CO_2_ uptake rate accelerated to maximum which was then maintained for approximately 48 hours before monotonically returning to the baseline as the culture entered stationary phase (**Figure 8G**). The same strong correlations between both cell growth and cumulative fixed carbon (**Figure 8H**) as well as volumetric cell growth and CO_2_ uptake rates were once again revealed.

**Figure 8.**
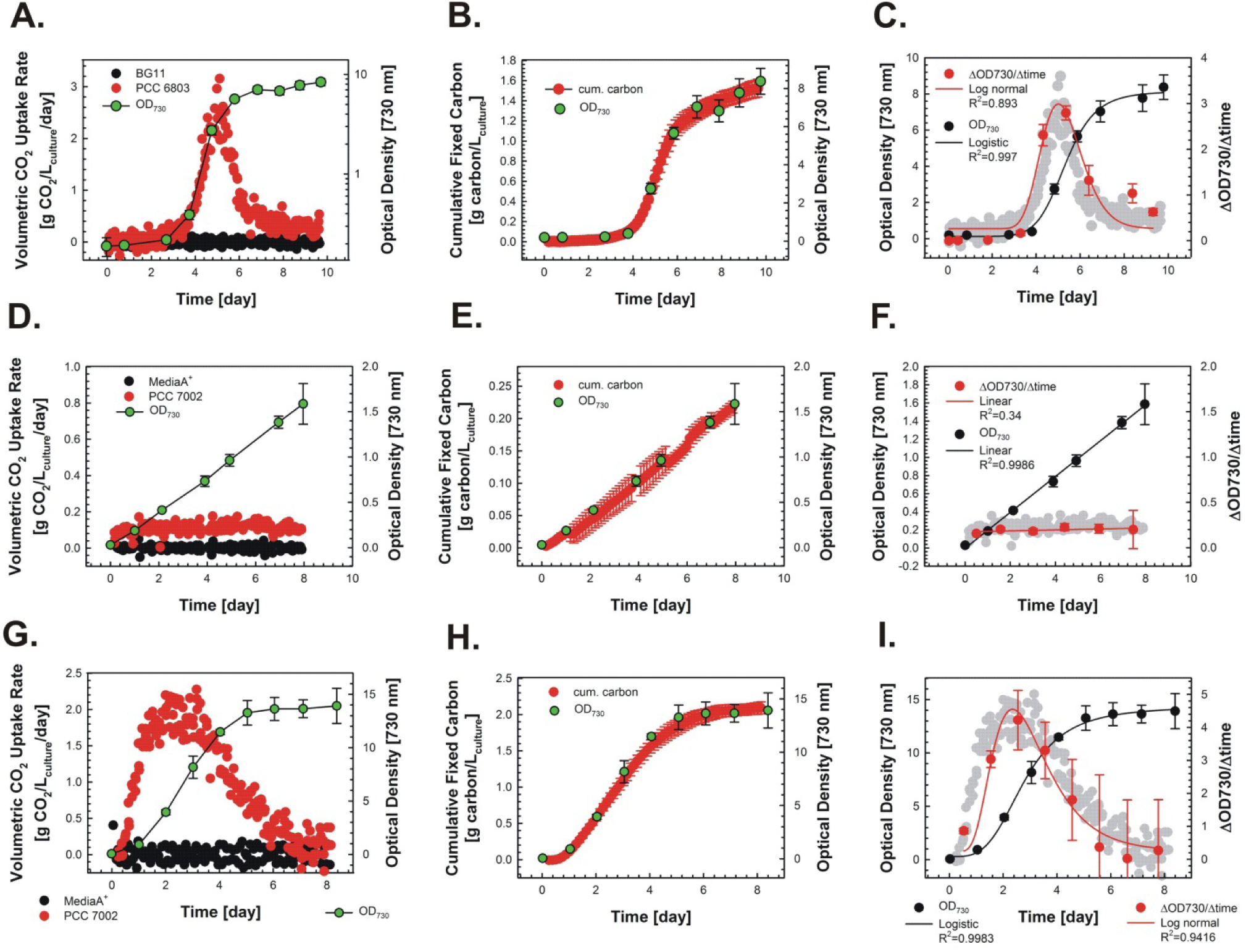
Correlating rates and levels of carbon fixation with those of cyanobacterial growth. **A-C)** *Synechocystis* sp. PCC 6803 growth at continuous illumination at 150 μE and 5,000 ppm (0.5%) CO_2_. **D-F)** *Synechococcus* sp. PCC 7002 growth at continuous illumination at 250 μE and air. **G-I)** *Synechococcus sp.* PCC 7002 growth at continuous illumination at 150 μE and 5,000 ppm (0.5%) CO_2_. **A,D,G)** Volumetric CO_2_ uptake rate and cell growth (OD_730_) determined instantaneously during batch culture. Superimposed is cell growth (OD_730_). **B,E,H)** Cumulative fixed carbon per volume of culture determined by integrating the data in A,D,G). Superimposed is cell growth (OD_730_). **C,F,I)** Cell growth (OD_730_; black circles) fit to the indicated best fit curve with the corresponding R-squared values. Volumetric cell growth (red circles) rate determined by taking the derivative of the growth curve with respect time and fit to a best-fit curve with the corresponding R-squared values. Volumetric CO_2_ uptake rate (gray circles) is underlaid behind the data growth rate data for comparison. Error bars represent the standard deviation from biological triplicates.

## 4. Conclusions

Here we report the construction, validation, and application of a low-cost, automated off-gas sampling system that accurately measures volumetric CO_2_ uptake rates in cyanobacterial cultures both instantaneously and with high temporal resolution. The unique datasets made available with this system have the potential to offer new and valuable insights regarding growth and CO_2_ fixation by cyanobacteria and other phototrophic microbes. Notably, the volumetric CO_2_ fixation rate is shown to be highly dynamic in nature, with steady-state growth (i.e., where the growth rate is both maximal and constant (Monod, 1949)) occurring for only a brief period during batch cultures of both PCC 7002 and PCC 6803. With the ability to understand the dynamic nature of CO_2_ fixation kinetics in cyanobacterial cultures, future studies aimed at metabolic flux analysis, ‘omics’ characterization, and metabolic engineering of diverse cyanobacteria will all be greatly enhanced. For example, in metabolic engineering applications, by understanding the magnitude of the maximum CO_2_ uptake rate one can then more accurately predict the theoretical maximum rate of product formation (assuming all carbon could be diverted away from biomass production). Meanwhile, since the maximum volumetric CO_2_ uptake rates occurs for just a brief period during batch cultures, the development of genetic strategies aimed at extending this most productive period will represent a new challenge for cyanobacterial synthetic biology.

Finally, it is noted that light-dependent CO_2_ gas exchange, or LCE (Spalding & Ogren, 1985), represents a similar technique to the one developed here. However, this technique: i) represents an *ex situ* method for measuring CO_2_ uptake rates since the cells are not examined under authentic batch conditions, and ii) involves time scales on the order of minutes, not days (Ohnishi et al., 2010; Yamano, Miura, & Fukuzawa, 2008). Our *in situ* approach is furthermore similar to other ‘CO_2_ balance’ methods employed by others (Shabestary et al., 2018; Shabestary et al., 2021), with the notable exception that our unique autosampler system allows for the facile acquisition of high-resolution datasets (i.e., 8 time points per day vs. just 1) and the monitoring of multiple cultures in parallel. With its low-cost design based on readily accessible parts, we envision this type of device becoming a commonly employed and valuable tool in cyanobacterial research.

