## Supplemental for "*In situ*, High-Resolution Quantification of CO_2_ Uptake Rates via Automated Off-Gas Analysis Illuminates Carbon Uptake Dynamics in Cyanobacterial Cultures"

### Supplemental Text

#### $CO_2$ and Flow Rate Measurements: Data Collection, Processing, and Analysis

To begin a 24 h recording period, the power to the solenoid-regulating Arduino, the serial monitor of the D6F-P Arduino program, and the recording function of the Vaisala infrared $CO_2$ gas meter were all initiated simultaneously. We used the time stamp feature of Vaisala infrared $CO_2$ gas meter and the D6F-P serial monitor to verify the synchronicity of the two devices. After 24 hours, sampling data from the Vaisala infrared $CO_2$ gas meter and the D6F-P serial monitor were copied into a spreadsheet. The timescale was adjusted to reflect the start of the experiment. For each tube, data collected during the final 5 min were averaged for both $CO_2$ and flow rate measurements. These resulting datasets (time [day], $CO_2$ [ppm], flow rate [L/min]) were copied into a second spreadsheet where the following calculations were performed.

To determine the mass delivery rate of $CO_2$ entering each culture tube, $\nu CO_{2(in)i}$, we used the equation:

$$\nu CO_{2(in)i} = \left[\frac{CO_{2(in)i}}{(1x10^6 \times R \times T)}\right] \times M_{r(CO2)} \times v_i$$

Where $CO_{2(in)i}$ is the $CO_2$ concentration ($ppm_v$) in the feed gas being delivered to the culture tube, as determined by sampling an empty tube in each loop. R is the universal gas constant (0.08206 L-atm/K-mol). T is temperature (K). $M_{r(CO2)}$ is the molecular weight of $CO_2$ (g/mol). $v_i$ is the flow rate (L/day) of gas to the culture tube, as determined by measuring the bubbling rate of the exiting off-gas using the D6F-P microflow sensor. $\nu CO_{2(in)i}$ is divided by the volume of culture to give the volumetric $CO_2$ delivery rate (g $CO_2/L_{culture}$/day). These data were averaged for each of the triplicate cultures over the course of an experiment to give a mean $CO_{2(in)i}$. To determine the mass rate of $CO_2$ exiting each culture tube, $\nu CO_{2(out)i}$, we used the equation:

$$vCO_{2(out)i} = \left[\frac{CO_{2(out)i}}{(1x10^6 \times R \times T)}\right] \times M_{r(CO2)} \times v_i$$

Where $CO_{2(out)i}$ is the $CO_2$ concentration ($ppm_v$) in the off-gas leaving the culture tube.

At this point, the ostensible $CO_2$ uptake value from each sample, $vCO_{2(in)i} - vCO_{2(out)i}$, does not account for a baseline $CO_2$ uptake value from the media blanks. We define in all our experiments that the mean $CO_2$ uptake rate of the blank media samples is zero over the course of an experiment. Therefore, to determine a media-adjusted volumetric $CO_2$ uptake rate we average the $CO_2$ uptake value from the media blanks and subtract that value from all samples. We call the mean $CO_2$ uptake rate from the blanks, $vCO_{2(media-avg)i}$. Thus, the volumetric $CO_2$ uptake rate is given by:

$$Volumetric\ CO2\ Uptake\ Rate_i = \left[vCO_{2(in)i} - vCO_{2(out)i}\right] - vCO_{2(media-avg)i}$$

To construct the $CO_2$ uptake curves, each of the $CO_2$ uptake rate data sets (time [day], volumetric $CO_2$ uptake rate [g $CO_2/L_{culture}$/day]) is combined and sorted from smallest to largest using a spreadsheet. Since the time values for each of the triplicate cultures are offset by 19 min, we reasoned that grouping the replicates in this manner gives a clearer and more accurate picture than averaging the three replicates. To determine the area under the curve we used the point-slope method of integration where x=time [day] and y=volumetric $CO_2$ uptake rate [g $CO_2/L_{culture}$/day]. The slope, $m_i$, is calculated by:

$$m_i = \frac{y_i - y_{i-1}}{x_i - x_{i-1}}$$

And the y-intercept, $b_i$, is calculated by:

$$b_i = y_i - (m_i \times x_i)$$

The area under the curve for each successive time point, $AUC_i$, is determined by:

$$AUC_i = \frac{\left[\left(\frac{m_i}{2}\right) \times \left(y_i^2 + b_i y_i\right)\right] - \left[\left(\frac{m_i}{2}\right) \times \left(x_{i-1}^2 + b_i x_{i-1}\right)\right]}{3.66}$$

A summation of the total area under the curve would represent the total amount of carbon fixed. However, the y-values at this point are in terms of g $CO_2$. Therefore, to convert to g carbon, integration value is divided by 3.66 which is the fold change in molecular weight between $CO_2$ and elemental carbon. A simple cumulative summation of all g-carbon adjusted AUCi's gives the total amount of carbon fixed.

**Mathematical fits**

All mathematical fits were generated using SigmaPlot v.11. The logistic fit used throughout is:

$$y = y_0 + \frac{a}{1 + \left(\frac{x}{x_0}\right)^b}$$

The Weibull function used in Figure 6 is:

$$y = a\left(\frac{c-1}{c}\right)^{\frac{1-c}{c}} \left[\frac{x - x_0}{b} + \left(\frac{c-1}{c}\right)^{\frac{1}{c}}\right]^{c-1} e^{-\left[\frac{x-x_0}{b} + \left(\frac{c-1}{c}\right)^{\frac{1}{c}}\right]^{c}} + \frac{c-1}{c}$$

The Lorentzian function used in Figure 7 is:

$$y = y_0 + \frac{a}{1 + \left(\frac{x - x_0}{b}\right)^2}$$

**Arduino programs**

Broadly speaking, the autosampler program activates a user-defined digital output pin on the Arduino microcontroller (Arduino Uno) for a user-defined period of time. Activating a given solenoid diverts the off gas from that sample into a collection manifold that is in line with a Vaisala infrared $CO_2$ gas meter and a D6F-P microflow sensor. The program is a loop that consists of 5 min of sampling the $CO_{2(in)}$ blank culture tube (to demarcate the beginning of a sampling loop), followed by 4 minutes of sampling a low-flow Nitrogen source (ca. 100 ml/min), followed by 15 minutes of sampling the blank culture tube (to determine $CO_{2(in)}$), followed by 4 minutes of sampling a low-flow Nitrogen source, followed by 15 minutes of sampling the first media blank (to set the baseline, to be averaged with the second media blank), followed by 4 minutes of sampling a low-flow Nitrogen source, followed by 15 minutes of sampling the first replicate of the first culture, followed by 4 minutes of sampling a low-flow Nitrogen source, followed by 15 minutes of sampling the second replicate of the first culture, followed by 4 minutes of sampling a low-flow Nitrogen source, followed by 15 minutes of sampling the third replicate of the first culture, followed by 4 minutes of sampling a low-flow Nitrogen source, followed by 15 minutes of sampling the second media blank, followed by 4 minutes of sampling a low-flow Nitrogen source, followed by 15 minutes of sampling the first replicate of the second culture, followed by 4 minutes of sampling a low-flow Nitrogen source, followed by 15 minutes of sampling the second replicate of the second culture, followed by 4 minutes of sampling a low-flow Nitrogen source, followed by sampling the third replicate of the second culture. The total loop time is equal to (n samples x 15 min) + (n+1 samples x 4 min) + 5 min.

*<u>Arduino Sampling Program:</u>*

```
Int solenoidPin10 = 10;
int solenoidPin9 = 9;
int solenoidPin8 = 8;
int solenoidPin7 = 7;
int solenoidPin6 = 6;
int solenoidPin5 = 5;
int solenoidPin4 = 4;
```

```
int solenoidPin3 = 3;
int solenoidPin2 = 2;
int solenoidPin1 = 1;
void setup()
{
  pinMode(solenoidPin1, OUTPUT);        //Set pin 1 as the output for solenoid 1
  pinMode(solenoidPin2, OUTPUT);       //Set pin 2 as the output for solenoid 2
  pinMode(solenoidPin3, OUTPUT);       //Set pin 3 as the output for solenoid 3
  pinMode(solenoidPin4, OUTPUT);        //etc
  pinMode(solenoidPin5, OUTPUT);
  pinMode(solenoidPin6, OUTPUT);
  pinMode(solenoidPin7, OUTPUT);
  pinMode(solenoidPin8, OUTPUT);
  pinMode(solenoidPin9, OUTPUT);
  pinMode(solenoidPin10, OUTPUT);
 }

void loop()
{
digitalWrite(solenoidPin1, HIGH);     //solenoid 1 ON (CO2-in reference)
delay(300000);                        //300000 milliseconds = 5 min (delineates a sample loop)
digitalWrite(solenoidPin1, LOW);      //solenoid 1 OFF

digitalWrite(solenoidPin10, HIGH);     //solenoid 10 ON (N2 purge)
delay(240000);                        // 4 min (3 minutes was too short, 5 min too long)
digitalWrite(solenoidPin10, LOW);      //solenoid 10 OFF

  digitalWrite(solenoidPin1, HIGH);     //solenoid 1 ON (CO2-in reference)
```

```
 delay(900000);                        // 15 min (for rate of CO2-in calculations)
 digitalWrite(solenoidPin1, LOW);      // solenoid 10 OFF

digitalWrite(solenoidPin10, HIGH);      //solenoid 10 ON (N2 purge)
delay(240000);                        // 4 min
digitalWrite(solenoidPin10, LOW);       //solenoid 10 OFF

digitalWrite(solenoidPin2, HIGH);      // solenoid 2 ON (blank media replicate 1)
delay(900000);                        // 15 minutes
digitalWrite(solenoidPin2, LOW);

digitalWrite(solenoidPin10, HIGH);      //solenoid 10 ON (N2 purge)
 delay(240000);                        // 4 min
digitalWrite(solenoidPin10, LOW);       //solenoid 10 OFF

digitalWrite(solenoidPin3, HIGH);      // solenoid 3 ON (culture A replicate 1)
delay(900000);                        // 15 minutes
digitalWrite(solenoidPin3, LOW);

digitalWrite(solenoidPin10, HIGH);      //solenoid 10 ON (N2 purge)
delay(240000);                        // 4 min
digitalWrite(solenoidPin10, LOW);       //solenoid 10 OFF

 digitalWrite(solenoidPin4, HIGH);      // solenoid 4 ON (culture A replicate 2)
 delay(900000);                        // 15 minutes
 digitalWrite(solenoidPin4, LOW);
```

```
digitalWrite(solenoidPin10, HIGH);     //solenoid 10 ON (N2 purge)
  delay(240000);                       // 4 min
  digitalWrite(solenoidPin10, LOW);      //solenoid 10 OFF

digitalWrite(solenoidPin5, HIGH);     // solenoid 5 ON (culture A replicate 3)
delay(900000);                       // 15 minutes
digitalWrite(solenoidPin5, LOW);

digitalWrite(solenoidPin10, HIGH);     //solenoid 10 ON (N2 purge)
delay(240000);                       // 4 min
digitalWrite(solenoidPin10, LOW);      //solenoid 10 OFF

digitalWrite(solenoidPin6, HIGH);     // solenoid 6 ON (blank media replicate 2)
delay(900000);                       // 15 minutes
digitalWrite(solenoidPin6, LOW);

digitalWrite(solenoidPin10, HIGH);     //solenoid 10 ON (N2 purge)
delay(240000);                       // 4 min
digitalWrite(solenoidPin10, LOW);      //solenoid 10 OFF

digitalWrite(solenoidPin7, HIGH);     // solenoid 7 ON (culture B replicate 1)
delay(900000);                       // 15 minutes
digitalWrite(solenoidPin7, LOW);

digitalWrite(solenoidPin10, HIGH);     //solenoid 10 ON (N2 purge)
delay(240000);                       // 4 min
digitalWrite(solenoidPin10, LOW);      //solenoid 10 OFF
```

```
digitalWrite(solenoidPin8, HIGH);      // solenoid 8 ON (culture B replicate 2)
  delay(900000);                        // 15 minutes
  digitalWrite(solenoidPin8, LOW);


digitalWrite(solenoidPin10, HIGH);     //solenoid 10 ON (N2 purge)
delay(240000);                         // 4 min
digitalWrite(solenoidPin10, LOW);      //solenoid 10 OFF

digitalWrite(solenoidPin9, HIGH);      // solenoid 9 ON (culture B replicate 3)
delay(900000);                         // 15 minutes
digitalWrite(solenoidPin9, LOW);

digitalWrite(solenoidPin10, HIGH);     //solenoid 10 ON (N2 purge)
delay(240000);                         // 4 min
digitalWrite(solenoidPin10, LOW);      //solenoid 10 OFF

}
```

*D6F-P flow rate program:*

```
const int sensorPin = A0;   // hook up the V_out of the D6F-P to Pin A0
const float baselineFlow = 0;
int sensorVal = 0;
void setup() {
  Serial.begin(9600);
  }
void loop() {
   int sensorVal = analogRead(sensorPin);
   float voltage = (sensorVal/1024.0) * 5;
  Serial.print("\t | \t Volts: ");
  Serial.print(voltage);
  float flow = ((voltage - .536)/.017); // to be determined empirically by users calibration curve
  Serial.print("\t | \t flow rate ml/min: ");
  Serial.println(flow);

 delay(59929);   // 1 min sampling time minus the ca. 71 milliseconds it takes to run through the loop
}
```

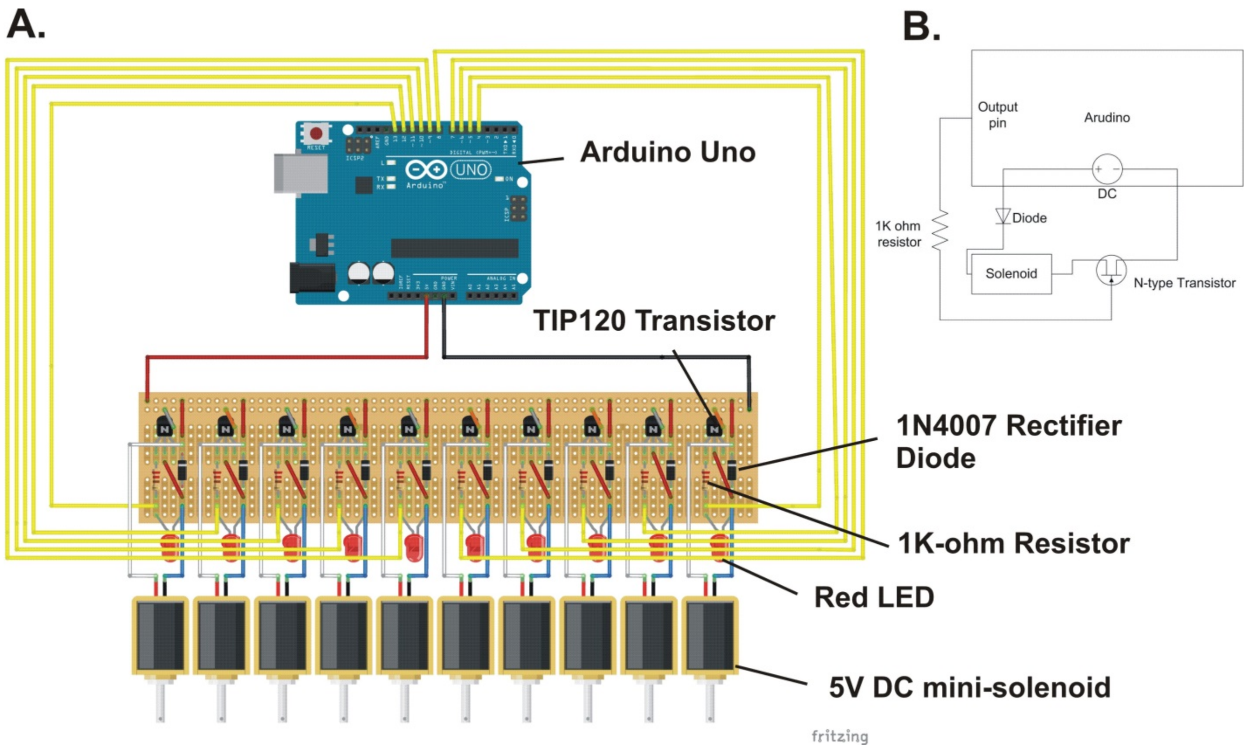


**Supplemental Figure 1. Circuit diagram of the autosampler. A)** Pictorial diagram (generated using the program Fritzing) with associated components as indicated. From left to right, the TIP120 transistor pins are: base, emitter, and collector. **B)** A generalized circuit diagram illustrating a single Arduino-controlled solenoid. Both the mini-solenoids and the Arduino microcontroller are powered by a 5V DC power supply.

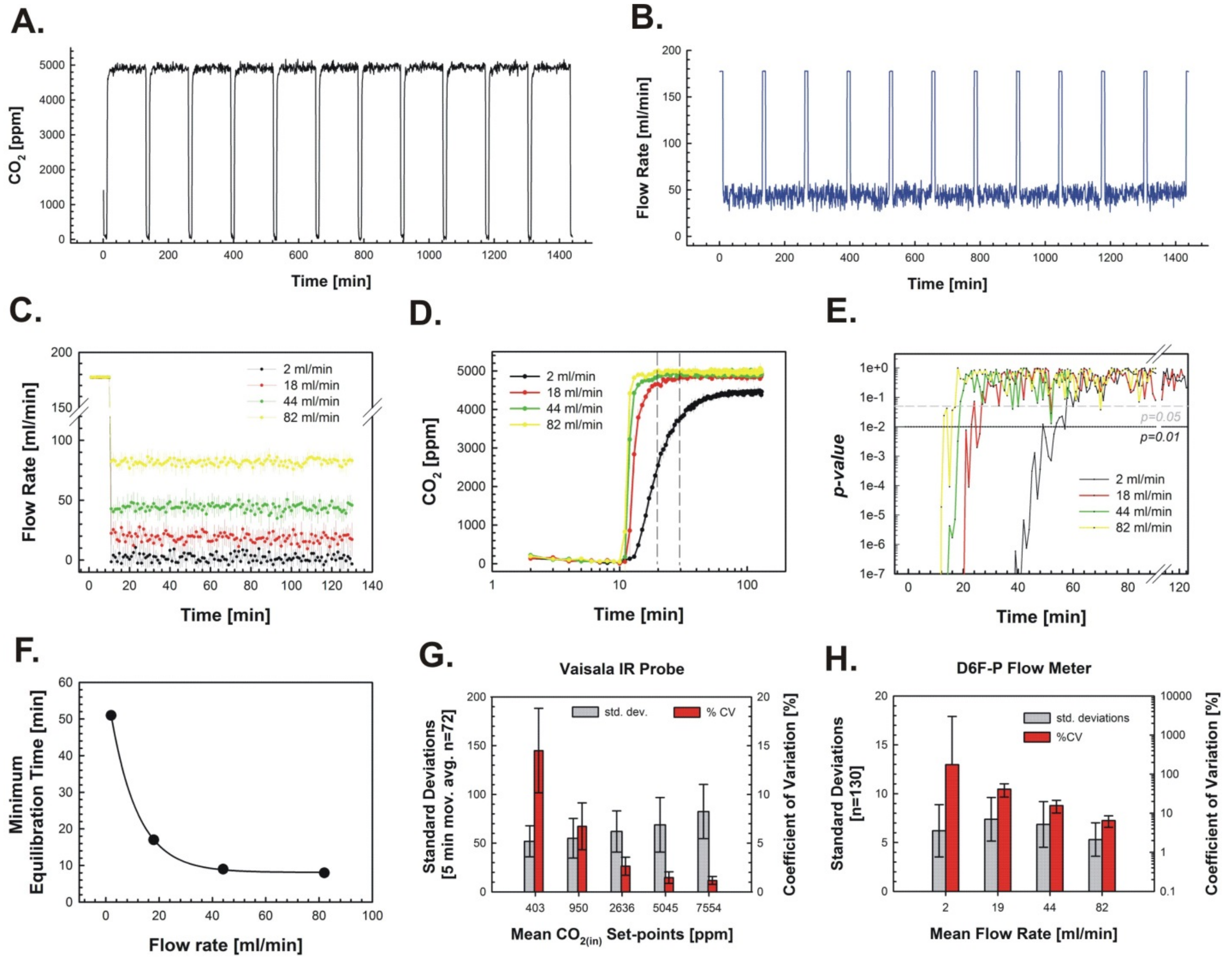


**Supplemental Figure 2. Determination of the minimum equilibration time for $CO_2$ probe and quantification of variance.** A solenoid sampling program was established to 1) sample $N_2$ for 10 min followed by 2) 120 min of continuous sampling of the off-gas from a 50 mL culture tube containing sterile BG-11 media bubbled with feed gas containing 5,000 ppm (0.5%) $CO_2$ at various flow rates. **A)** Example of off-gas $CO_2$ measurements obtained during repeated experiments at a flow rate of 44 mL/min. **B)** Example of flow rate measurements obtained during repeated experiments at a flow rate of 44 mL/min. **C)** Mean flow rates measurements (n=8) for each of the indicated settings. Error bars represent standard deviations. **D)** Mean off-gas $CO_2$ measurements (n=8) for each of the indicated settings. Error bars represent standard deviations. **E)** *p*-Values from a Student's *t*-test performed between the mean $CO_2$ measurement at each time point and the mean value during the final 30 min of equilibration. 'Minimum equilibration time' is defined as the time point corresponding to the first *p*-value greater than 0.05 that is then followed by 3 consecutive *p*-values greater than 0.05. **F)** Minimum equilibration time as a function of gas flow rate. **G)** Variance of the $CO_2$ gas probe at different $\boldsymbol{CO_{2(in)}}$ set points. **H)** Variance of the D6F-P micro flow meter at different gas flow rates.

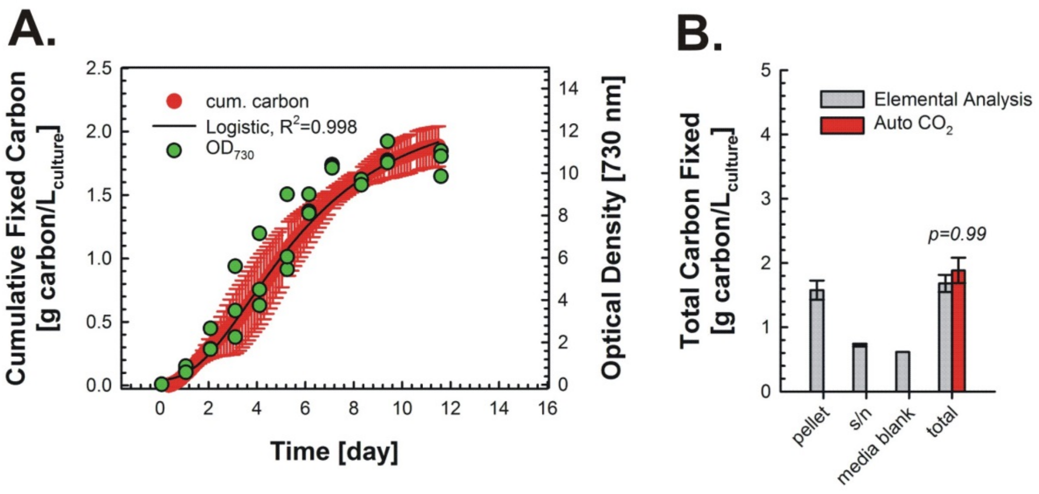


**Supplemental Figure 3. Validation of cumulative fixed carbon levels measured using the automated $CO_2$ off-gas sampling system.** Wild-type *Synechococcus* sp. PCC 7002 was grown in MediaA+ under continuous illumination at 150 μE and a feed gas containing 2,500 ppm (0.25%) $CO_2$. **A)** Cumulative fixed carbon, fitted with a sigmoidal function. Superimposed is cell growth ($OD_{730}$). **B)** Comparing total levels of carbon fixation by the culture as predicted via the off-gas sampling system versus measured via biomass elemental analysis. Total carbon analysis was performed using a PE 2400 CHN elemental analysis on the lyophilized cell pellet harvested at the end of the culture ('pellet') while a Shimadzu TOC-V analyzer was used for the supernatant ('s/n') and a MediaA+ blank. 'Total' was determined by a summation of the carbon content of both lyophilized cell pellets and media samples as described in the methods. Error bars represent the standard deviation from biological triplicates. (*p=0.99*, Student's *t*-test).

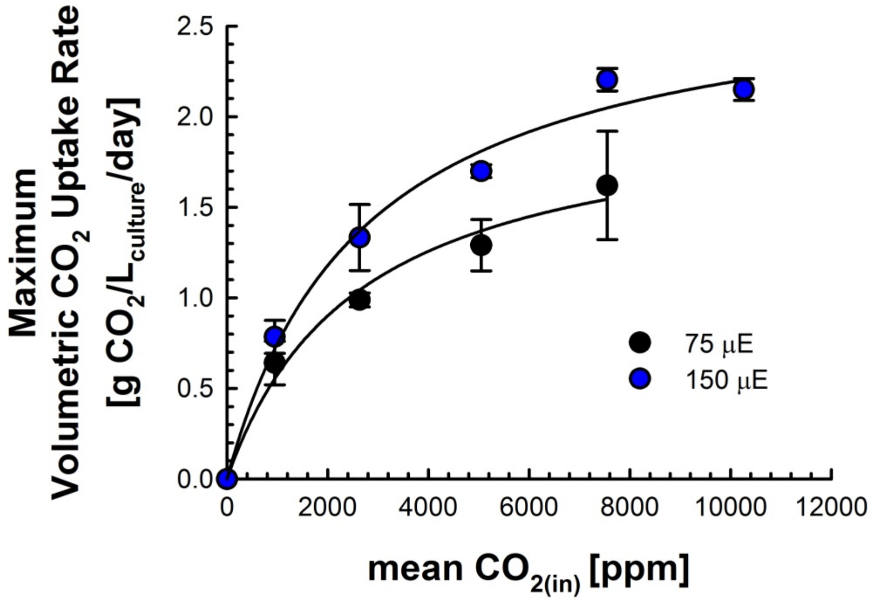


**Supplemental Figure 4. Comparing the effect of feed gas $CO_2$ concentration on $CO_2$ utilization efficiency. A)** Maximum $CO_2$ utilization efficiencies achieved when culturing PCC 7002 under continuous illumination at 75 or 150 μE and increasing feed gas $CO_{2(in)}$. All values calculated using the volumetric $CO_2$ uptake rate data from Figure 5. Error bars represent the standard deviation from biological triplicates.

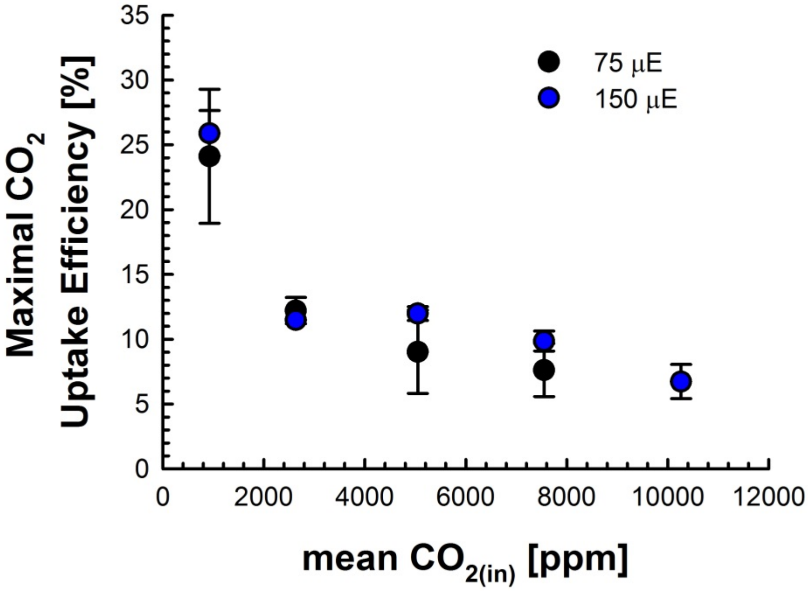


**Supplemental Figure 5. $CO_2$ uptake efficiency of PCC 7002 as a function of CO2(in).** Wild type PCC 7002 was grown in MediaA+ at the indicated $\boldsymbol{CO_{2(in)}}$ set point and continuous illumination at 75 or 150 μE. $CO_2$ uptake efficiency was determined as $\frac{vCO_{2(out)}}{vCO_{2(in)}} \times \mathbf{100}\%$. Error bars represent the standard deviations from the means of biological triplicates.

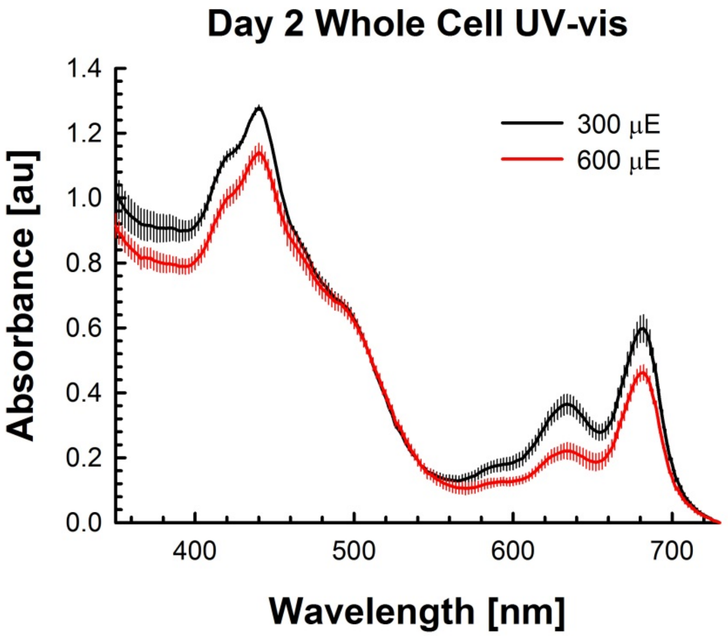


**Supplemental Figure 6. Whole cell UV-Vis analysis of PCC 7002 at elevated light intensities.** Wild type *Synechococcus sp.* PCC 7002 was grown under continuous illumination at 300 or 600 μE and 7,500 ppm (0.75%) $CO_2$ (collective data shown in Figure 7). On day 2, biomass has harvested from both cultures and whole cell UV-Vis spectroscopy was performed using a Tecan M1000 plate reader. Background absorbance was removed using a MediaA+ blank, after which absorbance readings were normalized by $OD_{730}$ followed by subtracting unity from each wavelength. Error bars represent the standard deviation from biological triplicates.
